# Why humans are stronger but not faster after isometric strength training: specific neural, not muscular, motor unit adaptations

**DOI:** 10.1101/2021.03.20.436242

**Authors:** A. Del Vecchio, A. Casolo, J. Dideriksen, P. Aagaard, F. Felici, D. Falla, D. Farina

## Abstract

While maximal force increases following short-term isometric strength training, the rate of force development (RFD) may remain relatively unaffected. The underlying neural and muscular mechanisms during rapid contractions after strength training are largely unknown. Since strength training increases the neural drive to muscles, it may be hypothesized that there are distinct neural or muscular adaptations determining the change in RFD independently of an increase in maximal force. Therefore, we examined motor unit population data during the rapid generation of force before and after four weeks of strength training. We observed that strength training did not change the RFD because it did not influence the number of motor units recruited per second or their initial discharge rate during rapid contractions. While strength training did not change motoneuron behaviour in the force increase phase of rapid contractions, it increased the discharge rate of motoneurons (by ∼4 spikes/s) when reaching the plateau phase (∼150 ms) of the rapid contractions, determining an increase in maximal force production. Computer simulations with a motor unit model that included neural and muscular properties, closely matched the experimental observations and demonstrated that the lack of change in RFD following training is primarily mediated by an unchanged maximal recruitment speed of motoneurons. These results demonstrate that maximal force and contraction speed are determined by different adaptations in motoneuron behaviour following strength training and indicate that increases in the recruitment speed of motoneurons are required to evoke training-induced increases in RFD.

## Introduction

Strength training leads to an increase in the neural output of the spinal cord during rapid (Van Cutsem *et al*., 1998*a*; Aagaard *et al*., 2002*b*; Vila-Chã *et al*., 2010; Tillin & Folland, 2014) and maximal voluntary contractions (Patten *et al*., 2001; Kamen & Knight, 2004). The increase in muscle force following strength training is mediated by an increase in discharge rates and reduced recruitment thresholds of motor units (Van Cutsem *et al*., 1998*a*; Del Vecchio *et al*., 2019*a*).

Despite an increase in maximal force, several studies adopting <12 weeks of isometric strength training have shown unchanged rate of force development (RFD) (Tillin & Folland, 2014; Balshaw *et al*., 2016). Moreover, when present, there is generally a large variability in the RFD responses after strength training (Del Balso & Cafarelli, 2007; Blazevich *et al*., 2020). The underlying neuromuscular mechanisms determining the changes in maximal force but not RFD after strength training are not fully known (Duchateau *et al*., 2005; Maffiuletti *et al*., 2016; Blazevich *et al*., 2020); both neural (Tillin & Folland, 2014; Balshaw *et al*., 2016) and muscular (Staron *et al*., 1994; Jürimäe *et al*., 1996; Seynnes *et al*., 2007*a*; Andersen *et al*., 2010; Tillin *et al*., 2012) adaptations have been discussed (Aagaard, 2003; Aagaard *et al*., 2020).

After short-term strength training, we previously observed that an increase in maximal force was associated to a compressed motor unit recruitment range and an increase in discharge rates during slow isometric ramp contractions (Del Vecchio *et al*., 2019*a*). Moreover, previous studies reported changes in muscle morphology with short-term strength training (Staron *et al*., 1994; Jürimäe *et al*., 1996; Seynnes *et al*., 2007*b*). Nonetheless, the muscle force twitch responses evoked by peripheral nerve stimulation are unchanged following short-term strength training (Van Cutsem *et al*., 1998*a*; Carroll *et al*., 2002; Nuzzo *et al*., 2017).

The main determinants of RFD are the recruitment speed and discharge rate of motoneurons (Desmedt & Godaux, 1978; Duchateau & Baudry, 2014; Del Vecchio *et al*., 2019*c*; Dideriksen *et al*., 2020). Because some studies have reported RFD to remain unchanged following strength training (Tillin & Folland, 2014; Balshaw *et al*., 2016), strength training presumably may not always influence the motor unit discharge timings (such as, recruitment speed, peak discharge rate during rapid contractions). In contrast, strength training mediates a more compressed motor unit recruitment and increase in motor unit discharge rates during slow contractions (Del Vecchio *et al*., 2019*a*), which should influence the rapid generation of force (Duchateau & Baudry, 2014; Del Vecchio *et al*., 2019*c*). Moreover, it has been proposed that the similar RFD after strength training reported in some studies, may be due to a combination of neural adaptations that tend to increase RFD and non-neural (muscular, tendinous, or myoelastic/myoelectrical components) adaptations that counteract these effects (Andersen *et al*., 2010; Balshaw *et al*., 2016; Blazevich *et al*., 2020).

Using robust methods for detecting motor unit activity during fast movements (Del Vecchio *et al*., 2019*c*), in this study we directly observed the behaviour of motor units during rapid contractions before and after strength training. The study was designed to investigate the adaptive plasticity of the motoneuron pool in response to short-term strength training, with an experimental focus on the adaptations in the output from the spinal cord during rapid contractions. The experimental results were supported with the results of numerical simulations.

We have previously reported the adaptations of the proposed training regime on the behaviour of the spinal motoneurons during slow force contractions (5% maximal voluntary force (MVC) per second (Del Vecchio *et al*., 2019*b*)), here we focus on fast movements, which consisted of contractions with rate of force developments as high as 600% MVC per second. Overall, the results clarify current conflicting evidence on changes in RFD with strength training and indicate that neural, not muscular, mechanisms are responsible for the inability to increase the RFD after isometric strength training.

## Material & Methods

### Participants

Twenty-eight healthy, non-smoking, young men volunteered to participate in this study, which conformed with the standards set by the *Declaration of Helsinki* and were approved by the Ethical Committee of the University of Rome ‘Foro Italico’ (approval no. 44680). Written informed consent was obtained from all participants prior to inclusion.

Exclusion criteria were age < 18 and > 30 years, current or previous history of any neuromuscular disorder and/or traumatic lower body injury and/or surgery, and regular involvement in any physical exercise program. Volunteers’ eligibility to the study and physical activity habits were assessed with a standard health survey and with the International Physical Activity Questionnaire (IPAQ, (Craig *et al*., 2003)), respectively, prior to their enrolment. Participants were recreationally active individuals, involved in light-to-moderate intensity physical activity up to a maximum of twice per week.

Participants were randomly assigned to either an intervention group (INT) or a control group (CON). Three participants dropped out for personal reasons (e.g. time demands), which resulted in a total of 25 participants (INT, n = 13; age, 23.9 ± 2.9 yr., weight, 74.1 ± 9.0 kg; height, 1.77 ± 0.08 m; CON, n = 12; age, 25.1 ± 2.9 yr. weight, 73.3 ± 8.0 kg; height, 1.78 ± 0.06 m) completing the study.

### Study overview

The characteristics of the strength training programme have been described in detail previously (Del Vecchio *et al*., 2019*a*; Casolo *et al*., 2019) and are briefly described here.

Participants attended the laboratory on 15 separate occasions over a 7-week period. After a familiarization session (*Session 1*), two duplicate main measurement sessions (baseline assessment, *Session 2;* post-intervention assessment, *Session 15*) were conducted four weeks apart. The familiarization session involved maximal, rapid and submaximal voluntary isometric ankle dorsiflexion with the dominant foot (self-reported). In the familiarization session, participants completed the same contractions of the main protocol, without recording of electromyography (EMG).

*Session 2* was carried out 3-to-5 days after *Session 1*, while *Session 15* occurred four weeks after *Session 2* (∼48-72 hours after *Session 14*). Each of the two-measurement sessions involved the concurrent recordings of ankle dorsiflexion force and myoelectrical activity of the tibialis anterior recorded with high-density surface electromyography (HDsEMG), whilst participants performed maximal, rapid and submaximal isometric contractions. *Session 3* to *14* were dedicated to the supervised isometric strength training intervention (3 sessions per week for 4 weeks), which involved unilateral (dominant) isometric rapid and sustained contractions of the ankle dorsiflexors (see *Training Protocol and* Fig. 1 B-C). Participants of both groups were asked not to change their physical activity habits or their diet across the 4-week intervention. Participants were instructed to abstain from strenuous physical exercise and caffeine consumption for 48 h and 24 h prior to each measurement sessions, respectively.

**Figure 1.**
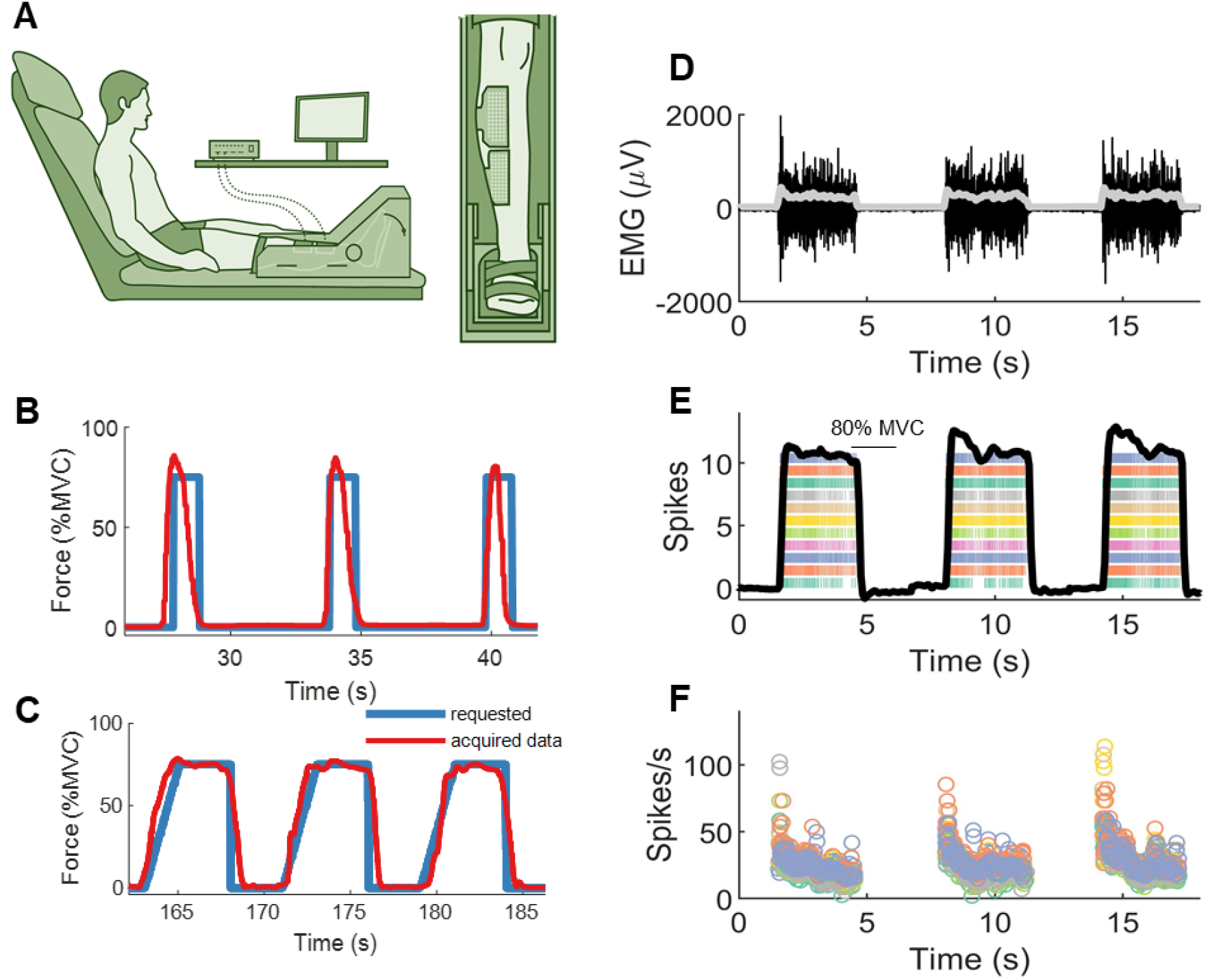
Motoneuron decomposition during rapid movements. **A**. The isometric dynamometer used for training and measurement sessions. **B**. Overview of the rapid contractions (4 sets x 10 repetitions) included in the training. Three contractions are shown for one representative subject (in blue, visual template with a horizontal cursor placed at 75% of MVC; in red, acquired force data). **C**. Overview of the sustained ramp contractions (3 sets x 10 repetitions) included in the training. Three contractions are shown for one representative subject (in blue, visual template characterized by a linear increase in force (37.5 MVC·s^-1^) up to a target level set at 75% of MVC and a 3 s plateau phase; in red, acquired force data). **D**. One bipolar EMG signal out of 64 acquired monopolar recordings during three isometric rapid force contractions for one representative subject. **E**. Colour-coded raster plots of the individual motor units identified from the tibialis anterior muscle of one representative subject during one measurement session (in black, acquired force trace in % of MVC. **F**. The instantaneous discharge rate for all the motor units that are shown in the raster plot panel (**E**).

### Experimental procedures

A standardized and progressive warm-up consisting of eight isometric contractions of dorsiflexion at different intensities of self-perceived maximal voluntary force (4 x 50%, 3 x 70%, 1 x 90%, with 15-30 s rest), and a series of rapid dorsiflexion contractions was performed at the beginning of each measurement session. For the rapid contractions, which were used to familiarize the participants with the development of maximal volitional force as rapidly as possible, participants were instructed to contract “as fast as possible” up to a target level displayed on a screen (∼75% of perceived maximum voluntary force), and to release immediately after the achievement of the peak force (Van Cutsem *et al*., 1998*b*; Del Vecchio *et al*., 2019*c*). With this protocol, we have observed a strong agreement in maximal RFD achieved during rapid contractions targeting a submaximal force level vs. intending to reach maximal contraction force (Del Vecchio *et al*., 2019*c*).

Following the warm-up, participants performed 3-to-4 maximal isometric voluntary contractions (MVCs), separated by 30 s of rest between trials. In each trial, participants were instructed to “pull as hard as possible” for 3-5 s, while being verbally encouraged to exceed the previously obtained force level, which was displayed on a monitor. The highest instantaneous force value recorded among the three-ankle dorsiflexion MVCs, i.e. the maximal voluntary isometric force, was used as a reference to determine the target forces for the submaximal trials.

Approximately 4 min after the completions of the MVCs, participants performed 12 rapid isometric contractions of ankle dorsiflexion (2 blocks of 6 repetitions each, with 20 s and 2 min of rest in between, respectively). The volunteers were instructed to contract “as fast and as hard as possible” in order to exceed a horizontal cursor displayed on a monitor, set at 75% of their maximal voluntary isometric force, and to maintain a constant force for 3 s at the target force level, before returning to baseline values. An auditory cue was provided prior to each rapid contraction, and participants were instructed to avoid any counter-movement or pre-tension. This type of contraction, characterized by an initial phase where the participants had to generate their maximum explosive force in the shortest period of time (i.e. maximal RFD), and followed by a 3 s plateau phase where the participants had to maintain a constant level of volitional force at the target.

### Training protocol

The training intervention comprised a total of 12 supervised sessions over 4 weeks. Each session lasted ∼30 min, and subsequent sessions were separated by 48-72 h. The training was performed in a laboratory setting on the same ankle ergometer (OT Bioelettronica, Turin, Italy) used for baseline and post-test assessments.

Following a standardized and progressive warm-up, characterized by five submaximal isometric contractions (2 x 50, 2 x 70, 1 x 90% of perceived MVC), participants performed ankle dorsiflexion MVCs (x 3) and a combination of rapid (4 sets x 10 repetitions, 1 min and 5 s of rest in between, respectively) and sustained ramp (3 sets x 10 repetitions, 2 min and 2 s of rest in between, respectively) isometric ankle dorsiflexion contractions with the dominant foot. Figure 1B-C shows an example of the slow and fast contractions performed during a training session. The MVCs were used as reference for the determination of submaximal contraction intensities. In the rapid contractions, participants were instructed to contract “as fast and as hard as possible” for ∼1 s in order to exceed a horizontal cursor set at their 75% of MVC, without any counter movement and/or pre-tension, and to release immediately thereafter. In the sustained ramp contractions, participants were instructed to match as precisely as possible a visual template, characterized by a linear increase in force at a constant rate (37.5 MVF·s^-1^) up to a target force level (75% of MVC), and to maintain a plateau for 3 s.

### Force recording

The familiarization and the main measurement sessions were completed with the same custom-built ankle ergometer (OT Bioelettronica, Turin, Italy), whose participant-specific configuration was established and registered during *Session 1* and precisely replicated thereafter. Participants were comfortably seated with their back against the seat back with their hip flexed to ∼120° (180° = anatomical position), their dominant knee extended to ∼180°, and the ankle positioned in ∼100° (90° = perpendicular to the tibia) of plantar flexion. The foot rested on an adjustable foot plate, in turn connected in series with a calibrated load cell (CCT Transducer S. A. S., Turin, Italy), which was positioned perpendicular to the plantar surface of the foot. The knee, ankle, and foot of the dominant limb were firmly strapped with Velcro straps (∼3 cm width), which were fastened above the patella, on the foot dorsum and on the distal portion of metatarsals, respectively. The contralateral leg (non-dominant) rested on the plinth to which the ankle ergometer was secured.

The analogue signal recorded by the load cell was amplified (x 200), A/D sampled at 2048 Hz with a 16-bit external analogue-to-digital converter (EMG-Quattrocento, OT Bioelettronica, Turin, Italy). All force and HDsEMG signals were synchronously acquired with the software OT Biolab, Ver. 2.0.6352.0 (OT Bioelettronica, Turin, Italy). Visual feedback of the force produced during each contraction was provided to participants on a pc-screen (visual gain, 88 x 88 pixs/%MVC) along with a display of the specific target force templates (LabVIEW, Ver. 8.0, National Instruments, Austin, TX, USA).

### Force analysis

The force signal was converted to newtons (N) and subsequently low-pass filtered. The offset of force was corrected to account for the effect of gravity. Contractions that showed pre-tension or counter-movement (baseline force ≥0.5 N in 150 ms prior to force onset) were excluded by visual inspection, and subsequently repeated. The force signal was lowpass filtered with a zero-lag 4^th^ order Butterworth filter with a cut-off frequency of 400 Hz. This large bandwidth guaranteed high accuracy when visually determining the onset of force (Tillin *et al*., 2013). The onset of force was identified visually using a validated methodology (Tillin *et al*., 2010). After the identification of the force onset we filtered the force signals with a 20 Hz low-pass zero-lag 4th-order Butterworth filter. This type of filter eliminates any spurious activities in the force time series guarantying a non-delayed force output in comparison to the original signal (Del Vecchio *et al*., 2018).

We extracted several parameters from the time-force curve. First, the absolute force value was extracted at different time points, 50, 100, 150, 200, 300 ms after force onset. The RFD (interval mean of the first derivative of the force signal) was also extracted in the same time windows (T_0-50_, T_0-100_, T_0-150_, T_0-200_ ms). In these time windows, we also extracted the Impulse, which corresponds to the time force integral. We either averaged these values across the three best contractions (i.e. showing the highest maximal RFD) or across contractions showing maximal RFD. Both the motor unit and force estimates were extracted in the same fashion from the motor unit model.

### EMG recording

HDsEMG signals were recorded from the tibialis anterior muscle of the dominant leg with two semi-disposal adhesive grids of 64 equally spaced electrodes (13 rows x 5 columns; gold-coated; 1 mm diameter; 8 mm inter-electrode distance; OT Bioelettronica, Turin, Italy). After skin preparation (shaving, gentle skin abrasion and cleansing with 70% ethanol), an experienced investigator identified the tibialis anterior muscle belly via palpation and marked its profile with a surgical pen. Two high-density electrode grids were positioned as described previously (Del Vecchio *et al*., 2019*a*; Casolo *et al*., 2019). Briefly, one adhesive grid was placed over the distal portion of tibialis anterior with the first four rows of electrodes on the identified innervation zone, and the first column aligned to the estimated anatomical direction of muscle fibres. The second adhesive grid was attached proximally to the first, in order to cover most of the muscle belly, thus maximizing the probability of sampling from as many independent sources as possible (motor unit action potentials). Disposable biadhesive foam layers (SpesMedica, Battipaglia, Italy), covered with holes filled with conductive paste, were used to attach the grids to the skin overlying the muscle and to optimize the skin-to-electrode contact.

The main ground electrode (strap electrode, dampened with water) was placed in proximity of the styloid process of the ulna of the tested side. Two reference electrodes (one for each electrode grid) were placed on the tuberosity of the tibia and on the medial malleolus of the tested limb. To replicate electrode positioning before and after the training period, the exact positioning of the two high-density grids were marked on the participants’ skin at the end of the baseline session and of each training session using tattoo ink.

All HDsEMG signals were recorded in monopolar configuration, sampled at 2048 Hz, amplified (x 150), band-pass filtered (10-500 Hz) at source, and converted to digital data with a multichannel amplifier with 16-bit resolution (EMG-Quattrocento, OT Bioelettronica, Turin, Italy), before being stored for offline analyses (MATLAB 2020, Mathworks Inc., Natick, MA, USA).

### EMG analysis

The raw HDsEMG signals recorded during submaximal isometric trapezoidal contractions (35-50-70% MVC) were digitally band-pass filtered (20-500 Hz, 2^nd^ order Butterworth) before being decomposed offline with a validated decomposition algorithm based on convolutive blind source separation method (Holobar & Zazula, 2007). The algorithm allows a highly reliable identification of motor unit discharge timings over a broad range of voluntary forces, including rapid contractions (Holobar *et al*., 2014; Del Vecchio *et al*., 2019*c*). Once the spike instants were identified by blind source separation, we performed a series of reinforcements and validity assessments of the motor unit spike trains, as described previously (Del Vecchio *et al*., 2019*c*). Briefly, the HDsEMG signals consist of convolutive mixtures of motor neuron spike trains. The relatively large dimension of the electrode grid allows for an accurate spatial sampling of the waveforms of the motor unit action potentials. The discharge timings of single motor units were converted to binary spike trains (0 = no firing; 1 = firing) and manually inspected by an experienced investigator. The accuracy of the decomposition procedure was ensured by retaining only those motor units characterized by a pulse-to-noise ratio (PNR) > 30 dB (Holobar *et al*., 2014). Moreover, only motor units that showed a consistently repeatable discharge pattern were considered for further analysis.

From all identified motor units, we computed the instantaneous discharge rate, cumulative spike trains, and the recruitment speed of motoneurons. The recruitment speed of motoneurons corresponds to the average time interval between the recruitment of the population of identified motoneurons. After the computation of the recruitment thresholds it was possible to sort the specific motor unit by recruitment order. Therefore, the average of the first time-derivative of this vector corresponded to an estimate of the recruitment speed of the motoneuron pool.

The discharge rate at recruitment corresponded to the average instantaneous discharge rate of the first three interspike (two values) intervals and during the plateau phase (average of 10 spikes, yielding 9 values), at 300 ms from contraction onset. We also computed the average discharge rate from the cumulative spike trains. For this purpose, a moving average of 35 ms with an overlap of 1 ms beginning during the first motor unit spike was obtained from the cumulative spike train. During each window, the total number of firings was divided by the number of active motor units and the window length to yield the average motor unit discharge rate. The instantaneous discharge rate was averaged across the pool of identified motor units and also stored for each motor unit and contraction intensity. For statistical purposes, we only compared the two cohorts with the motor unit averaged data. However, we also present in the results and figures the motor unit discharge characteristics for the full population of identified units.

### Motor unit model

The motor unit model has been described and verified previously (Dideriksen *et al*., 2020). The model adopted motor neuron discharge characteristics from experimental data and simulated corresponding isometric force output using a previously published model (Fuglevand *et al*., 1993). In the model of the single motor unit, discharge characteristics were determined based on three variables: the initial discharge rate of motoneurons, the discharge rate at plateau of force and the recruitment interval (time from recruitment of the first to the last motor unit). The discharge rates declined exponentially over a 250 ms period from the initial to the plateau rate. Within the recruitment interval, motor units were recruited according to size principle with a timing determined based on their assigned motor unit twitch force amplitudes. The recruited motor units were used to model the twitch of the motor units. We performed two sets of simulations. *Simulation 1* generated a number of contractions that matched the experimental data. For these simulations we used the discharge rate values before and after the intervention for the 12 participants in the control group and 13 in the intervention group. For *Simulation 2*, we generated 400 contractions from a random gaussian distribution of motor unit discharge characteristics obtained from the average and standard deviation of the experimental values. For both simulations, the muscle gain (motor unit twitch force amplitudes) was the same. Therefore, any of the potential changes in force output could solely be attributed to changes in motoneuron output.

### Statistical analysis

The Shapiro-Wilk test was used to assess the normality of distribution of data. The majority of the variables included in the analysis did not deviate from normal distribution. Changes in MVC, and neural properties (i.e. initial motor unit discharge rate, at plateau and motor unit recruitment speed) during the rapid isometric contractions were examined with multiple two-way repeated measure analysis of variance (RM ANOVA), where independent variables were time (before vs. after) and group (intervention, INT vs. controls, CON). When significant interactions were found, multiple comparisons were adjusted with Bonferroni correction. Changes in force variables extracted from the time-force curve (i.e. absolute and normalized RFD, absolute and normalized Impulse) were investigated with multiple three-way RM ANOVAs, where independent variables were time, group, and time window (T_0-50_, T_0-100_, T_0-150_, T_0-200_ ms). All statistical analyses were performed using the software GraphPad Prism Ver. 8.0.2 (GraphPad Software, San Diego, California, USA). The significance level was set at α < 0.05 for all tests. In the text, data are reported as mean ± SD.

## Results

After four weeks of strength training there was a significant increase (+12%) in maximal voluntary isometric force produced by the ankle dorsiflexor muscles (interaction: time x group, *P* = 0.006; INT: PRE: 289.4 ± 63.9 vs. POST: 328.9 ± 61.1 N; Bonferroni adjusted *P* < 0.001), whereas no changes were observed for the CON group (PRE: 299.2 ± 40.6 vs. POST: 304.2 ± 35.7 N).

There were no changes in either absolute or normalized (with respect to maximal force) RFD or impulse (Fig. 2). Specifically, absolute RFD obtained at the different time intervals remained unchanged after training (interaction: time x group x window, *P* = 0.9103; Bonferroni adjusted *P* > 0.05 in all cases; Fig. 2A). Similarly, RFD normalized to peak MVC force did not differ before and after training at the different time points (interaction: time x group x window, *P* = 0.738, Bonferroni adjusted *P* > 0.05, in all cases). The impulse (time force integral) assessed at different time intervals (T_0-50_, T_0-100_, T_0-150_, T_0-200_) in both absolute (interaction: time x group x window, *P* = 0.995) and normalized (interaction: time x group x window, *P* = 0.629) values also remained unaltered across the 4-week intervention period; Bonferroni adjusted *P* > 0.05, in all cases (Fig 2B).

**Figure 2.**
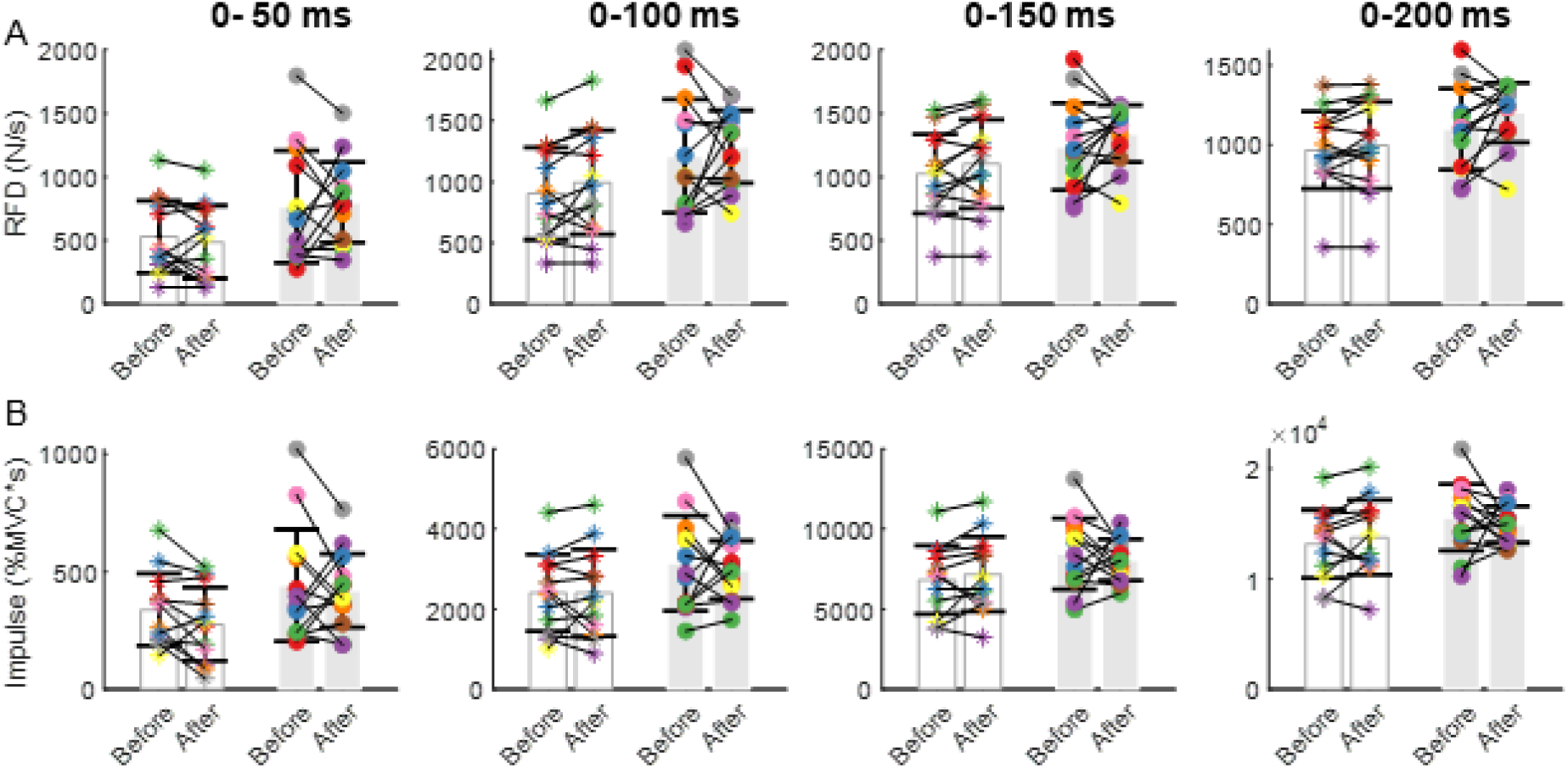
Estimates of ankle-dorsiflexor isometric contractile RFD extracted from the time-force curve before and after the 4-week intervention period (open bars: controls, grey bars: strength training). **A**. Absolute RFD (Newtons per second) in the different time windows (T_0-50_, T_0-100_, T_0-150_, T_0-200_ ms) from contraction onset. **B**. The integral of the normalized force-time curve (maximal voluntary force contraction times the window length in seconds) in the different time windows (T_0-50_, T_0-100_, T_0-150_, T_0-200_ ms) from contraction onset.

Because the primary determinants of RFD are known to be the recruitment speed of motoneurons and discharge rate of motor units during the first milliseconds of contraction, we investigated the motor unit behaviour in short time windows.

Figure 3 shows the instantaneous discharge rate at recruitment and plateau for all the motor units identified across the different contractions (participants are color coded). Strength training did not change the initial discharge rate of motoneurons (first three interspike intervals) during rapid force contractions (PRE: 54.75 ± 12.89 spikes/s; POST to 57.54 ± 9.02 spikes/s; *P* = 0.620; Fig. 3 A-C). Conversely, the discharge rate at the plateau of force (300 ms) increased significantly (12 out of 13 individuals, PRE: 25.38 ± 3.37 spike/s; POST: 27.92 ± 3.51 spike/s; *P* = 0.001; Fig. 3 B-D).

**Figure 3.**
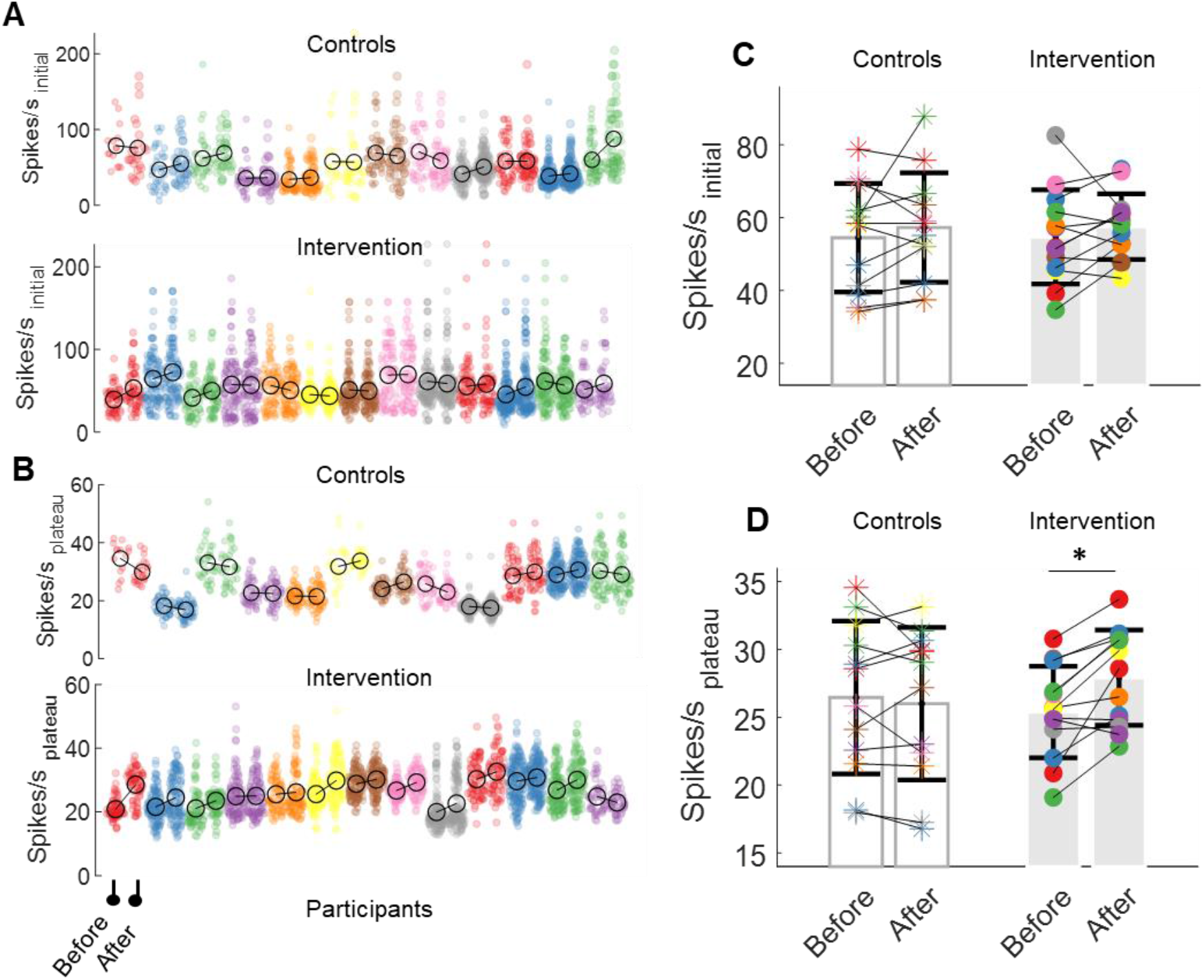
**A**. The average instantaneous discharge rate obtained from the first three spike intervals (Spikes/s initial, pps) for all the motor units that were identified for each subject during the rapid isometric contractions, in the controls (open bars) and intervention group (grey bars) before and after the four-week intervention period. Participants are color-coded. **B**. The instantaneous discharge rate averaged at the plateau of the rapid isometric contraction (10 spikes). Note that strength training resulted in a higher motor unit average discharge rate at the plateau for 12/13 participants but this was not sufficient to elicit significant changes to the RFD. **C**. The average motor unit discharge rate during the initial phase of the rapid contraction, single participants are color coded. **D**. The average motor unit discharge rate measured during the plateau of the rapid contractions was significantly higher (*P*= 0.001) after four-weeks of strength training (grey bars) whilst was unaltered in controls (open bars).

The estimated motoneuron recruitment speed (the time-derivative of recruitment intervals) was not significantly different before and after training (PRE: 49.6 ± 24.2; POST: 50.4 ± 20.4 motor units/ms; interaction: time x group; *P* = 0.860; Fig. 4).

**Figure 4.**
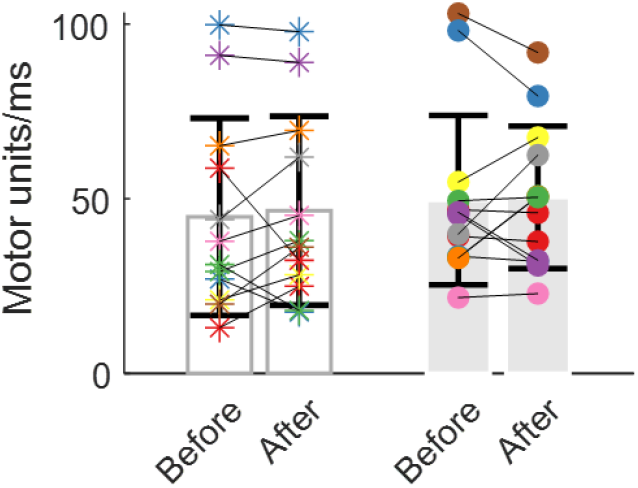
During rapid contractions, motor units are recruited separated by very brief time intervals. The recruitment speed of the population of identified motoneurons was calculated by computing the derivative of the recruitment intervals. Strength training (grey bars) did not change how fast motoneurons were recruited.

### Motor unit adaptations to strength training: Computer simulations

We have previously demonstrated with a motor unit model that the primary determinant of RFD is the recruitment speed of motoneurons (Dideriksen *et al*., 2020). Specifically, it was estimated that the increase of recruitment speed of motoneurons mediates a four-fold greater change in RFD than the effect of increasing the initial discharge rate and more than five-fold greater than increasing the chance of doublets (inter-spike intervals < 3 ms). The recruitment speed is also of greater importance when compared to the muscle characteristics (i.e. individual muscle fiber RFD), having an effect six-fold greater than twitch contraction times (Dideriksen *et al*., 2020).

We used a realistic motor unit force model responding to motoneuron firings mediated by strength training (Fuglevand *et al*., 1993; Dideriksen *et al*., 2020). Our aim was to study the relations between the training-induced change in motoneuron behaviour and the resulting changes in muscle force. Specifically, we assessed whether motoneuron discharge characteristics predicted RFD estimates and whether the increase in discharge rate would explain the change in maximal voluntary isometric force without a change in RFD. Two simulations were performed, one matching the experimental data (*Simulation 1*) and another comprising 400 permutated simulations obtained from a random gaussian distribution within the observed variability range (distributed across the mean and standard deviation of the population) of the observed motor unit parameters (*Simulation 2*). Figure 5A summarises the simulation conditions.

**Figure 5.**
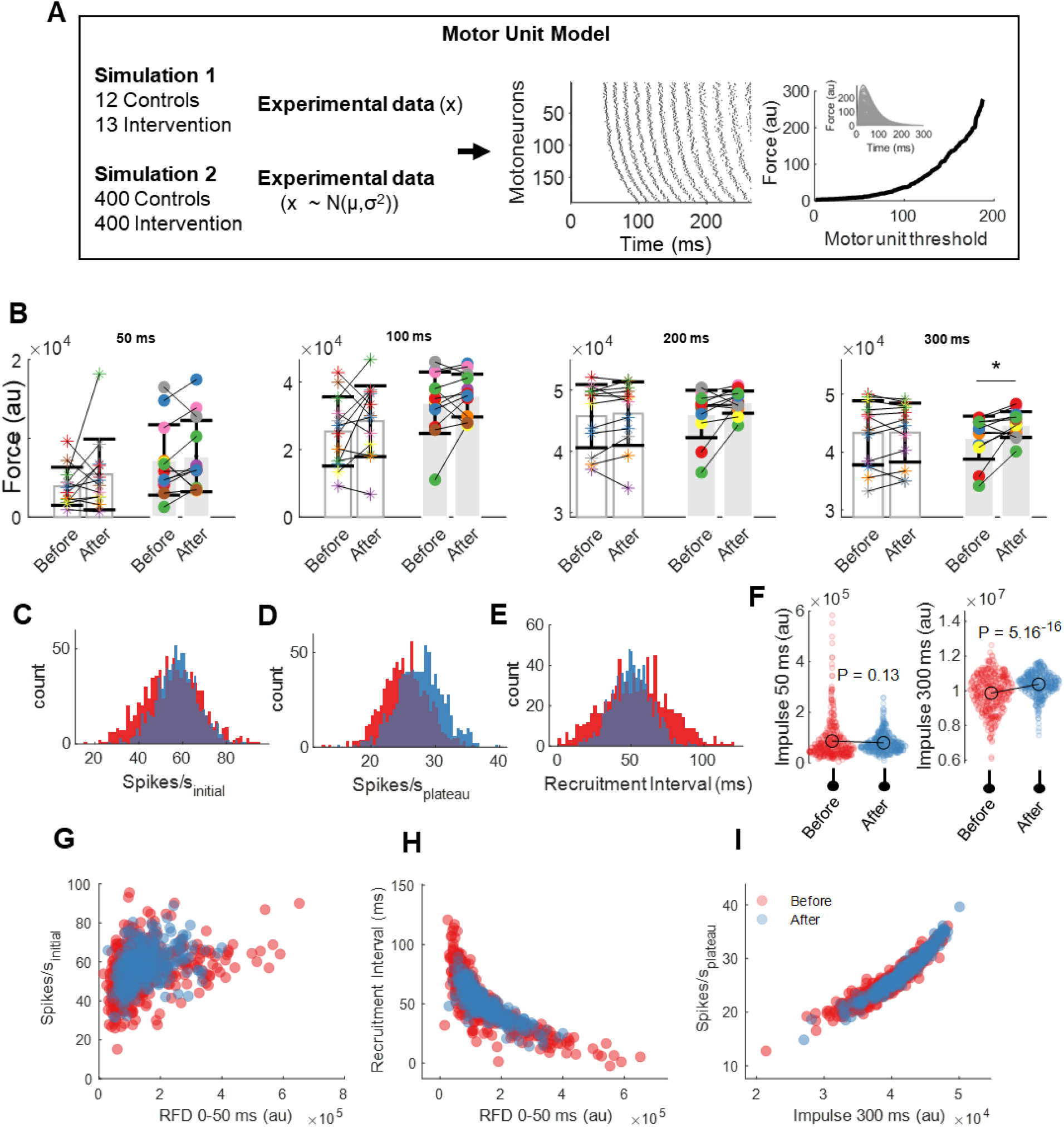
Simulation of rapid muscle force development with a motor unit model. **A**. Implementation of the simulation framework. Two simulations were adopted, the first simulation matched the experimental motor unit data at the individual subject level. In a second simulation we used the average and standard deviation of the various motor unit parameters to obtain 400 varying time-force curves. **B**. The force value at the individual subject level (colour-coded) at different time points (open bars: controls, grey bars: strength training; *P<0.001). **C**. Distribution of the random gaussian distribution in motor unit discharge rates (spike/s) in the initial phase of contraction (red bars indicate before and blue bars after the intervention). **D**. Spike/s distribution at the plateau of contraction. **E**. Distribution of recruitment intervals. Note the large variability in the post condition, matching the experimental variability in the recruitment interval data. **F**. The impulse (time * force) at 50 ms and 300 ms from force onset for the four hundred contractions, before and after training. **G**. The relation between early RFD and the initial motor unit discharge rate. **H**. Non-linear relation between early RFD and the recruitment interval. Note the significant increase in discharge rate with smaller recruitment intervals. **I**. Association between the cumulated impulse at 300 ms and the average discharge rate at the plateau of contraction. The Impulse a 300 ms is strongly correlated to maximal voluntary force.

The results from *Simulation 1* (12 controls and 13 from the intervention group) showed very similar force and RFD data as in the experimental conditions (Fig. 5B). The modelled force values at 300 ms from force onset were significantly higher (P<0.001) because of the higher discharge rate. Because the force values after 300 ms are correlated to maximal voluntary force values (Andersen *et al*., 2010; Folland *et al*., 2014), these results are well in agreement with our previous observations that the increase in maximal muscle force after four weeks of strength training is mediated by an increase in the discharge rate at the plateau phase but not at recruitment during slow force contractions (Del Vecchio *et al*., 2019*a*).

The motor unit discharge parameters that were used for the 400 simulations are shown in Fig 5C-I. The random sampling of the motor unit discharge characteristics resulted in a similar early rapid development of force and an increase in the force at the plateau of contraction (Fig. 5F). Interestingly, the change in discharge rate had a significant lower impact on the early RFD (Fig. 5G) than the recruitment speed of motoneurons, which showed a strong nonlinear dependence of early RFD on the recruitment interval of motor units (Fig. 5H).

One important aspect from these simulations is that in order for strength training to have an impact on early-phase RFD, the recruitment of motoneurons should happen in very small time intervals, i.e. at high recruitment speed. Indeed, the significant non-linear relation between early RFD and recruitment speed (Fig. 5H) was not dependent on the initial discharge rate (random gaussian distribution). On the contrary, variations in the initial discharge rate alone did not explain potential changes in RFD (Fig. 5G). However, at the plateau of force, when most motor units are fully recruited (∼300 ms), the increase in discharge rate was positively associated to maximal force (Fig. 5F). Indeed, there was also a significant relation between the impulse at 300 ms and the discharge rate (spikes/s plateau, Fig. 5I).

Collectively these simulations demonstrate the strong association between motor unit behaviour and muscle force production. They also show that recruitment speed is the main determinant of RFD, which explains why RFD did not change following training even in individuals for whom discharge rate increased during the early phase of the contraction (7/13 participants).

Figure 6 shows the time-force curve averaged in running 35-ms time intervals (overlap of 1 ms) across participants before and after training. In the same time interval, we also extracted the average motor unit discharge rate, obtained from the cumulative spike trains (Fig. 6B).

**Figure 6.**
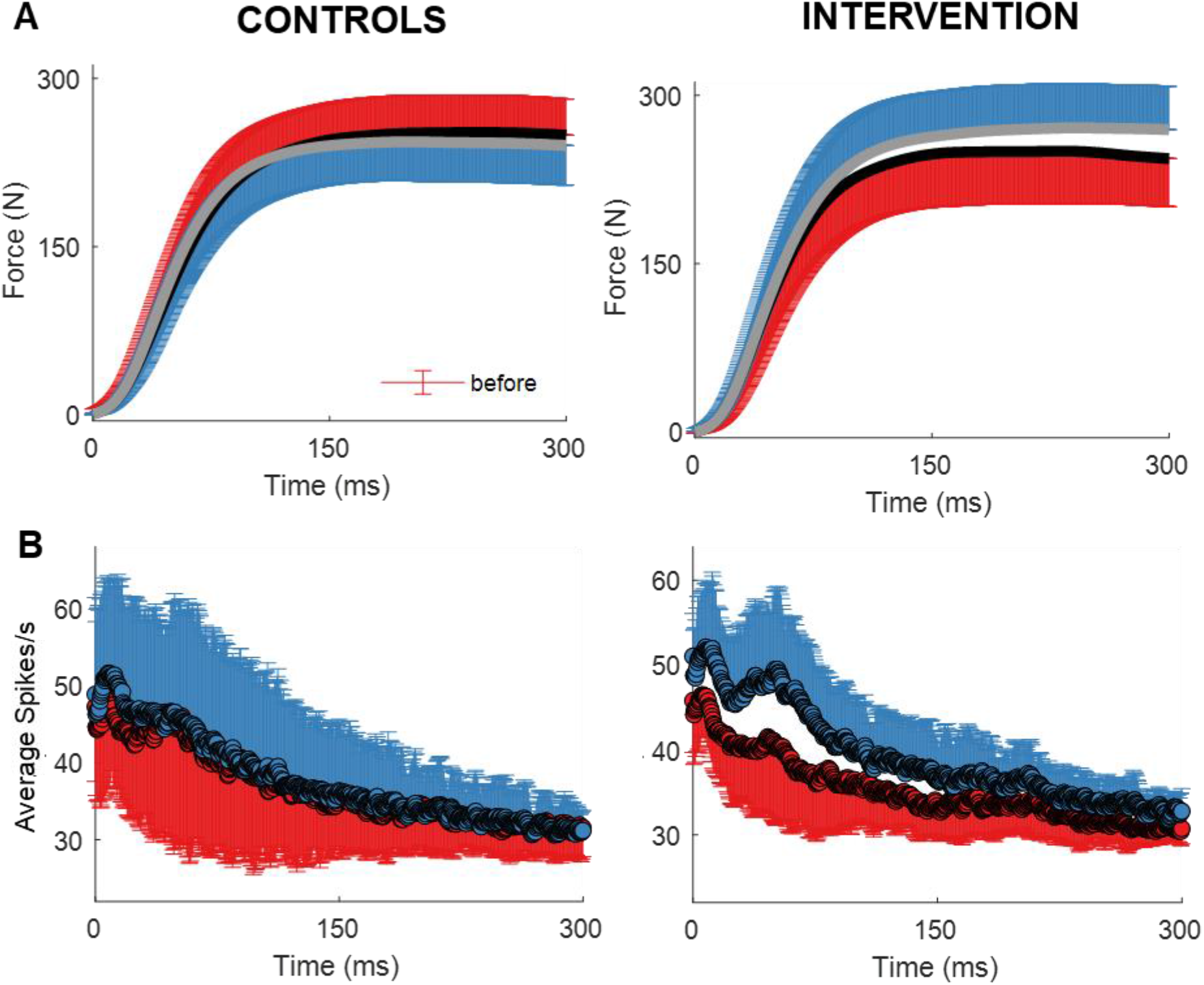
A The time-force curve averaged across all participants for the control and intervention group (group mean ±SD, red: before, blue: after the four-week intervention period, respectively). **B**. The average motor unit discharge rate (group mean ±SD) obtained in 35 ms time windows with an overlap of 1 ms.

## Discussion

We demonstrated with experimental data and computer simulations, that four weeks of isometric strength training of the dorsiflexors led to an increase in maximal muscle force production but did not influence the rate of force generation, as a result of specific motoneuron adaptations. We observed that strength training increased the discharge rate of motor units only in the later phase (+200 ms) of rising muscle force. In contrast, the recruitment speed and discharge rate of motoneurons did not change during the initial phase of the explosive contractions (<100 ms). During slow isometric contractions, motor unit recruitment thresholds decreased and motor unit discharge rate increased at the plateau phase (Del Vecchio *et al*., 2019*a*). These findings demonstrate that strength training elicits distinct motoneuron adaptations in the early vs later phases of rapid muscle contractions, so that maximal force increases without changes in RFD. The simulation model demonstrated that the neural adaptations alone could explain the observed divergent changes in force characteristics (maximal force and RFD) with strength training.

Previous studies have shown that in the absence of rapid muscle contractions, short-term isometric strength training may lead to an increase in the maximal force capacity of a muscle but with small or no effects on the RFD (Tillin & Folland, 2014; Balshaw *et al*., 2016). The underlying physiological processes governing these different responses were largely unknown. Some authors speculated that intrinsic changes within the muscle (i.e. sub-fiber type shifts from IIX to IIA) could negatively influence RFD (Andersen & Aagaard, 2010) whilst other studies showed a predominant neural basis for the differential training response (Tillin & Folland, 2014).

Interestingly, our training intervention increased maximal motor unit discharge rates during slow ramp contractions (at a fixed relative force intensity) (Del Vecchio *et al*., 2019*a*) as well as during the late, but not the early phase of rapid feedforward contractions. During rapid isometric contractions, motoneurons begin to discharge at high rates that determine the RFD (Desmedt & Godaux, 1977; Van Cutsem *et al*., 1998*b*; Duchateau & Enoka, 2011). The different behaviour of motoneurons after training as a function of time during a contraction may be due to divergent effects of strength training on afferent and supraspinal inputs. For example, one of the underlying mechanisms for the observed changes in force could be an increase in magnitude of excitatory Ia afferent input. Accordingly, previous studies have reported an increase in H-reflex and V-wave responses after strength training (Aagaard *et al*., 2002*b*; Duclay *et al*., 2008). However, while strength training may increase the afferent synaptic input to motoneurons, the time taken for the afferent volleys are too long to mediate faster RFD in the early phase of contraction (0-50 ms). Therefore, the increase in RFD with speed training may be determined mainly by an increase in the strength of supraspinal input.

Motoneuron pools during the fast and forceful development of force receive input from different descending pathways, such as the brainstem and cortex. Therefore, circuitries that determine speed and maximal force might be separated at the supraspinal level. Comparisons between training with rapid versus sustained contractions showed that if the exercise is performed with fast movements, the increase in RFD was achieved exclusively in those that trained with rapid contractions (∼14% gain) (Tillin & Folland, 2014; Balshaw *et al*., 2016). Indeed, it has been systematically reported that EMG amplitude increases following training with isometric rapid contractions while it does not change during rapid contractions following maximal strength training (Tillin & Folland, 2014; Balshaw *et al*., 2016). These findings indicate a neural basis mediating the inability to increase the RFD of human skeletal muscle. However, it is important to point out that when strength training is performed with dynamic movements, it results in changes in EMG and RFD (Aagaard *et al*., 2002*a*) and motor unit firing rates (Van Cutsem *et al*., 1998*a*). Our training intervention was very similar to the one adopted by (Tillin & Folland, 2014; Balshaw *et al*., 2016) during isometric knee extension. The time-force data was similar after the isometric protocol (Tillin & Folland, 2014; Balshaw *et al*., 2016) suggesting that these results may also be generalized to other joint compartments.

We used computer simulations to interpret and extend the experimental results. The muscle responses were modelled with a motor unit model with fixed motor unit twitches. This analysis allows to model the firing characteristics of the motoneurons and observe the responses of the time-force signals. The firing characteristics of the motoneurons were modelled according to the experimental data. The simulations demonstrated a clear association of motor unit recruitment speed with force, and that the observed variability in initial discharge rate of motoneurons is not a determinant of potential changes in muscle force. These results show that the RFD of a muscle is highly related to the recruitment speed of motoneurons and that future training intervention should focus on enhancing the recruitment speed of motoneurons.

In conclusion, we presented the behaviour of human motoneurons during rapid contractions after four-weeks of strength training. Our results demonstrate for the first time that strength training does not elicit changes in rate of force generation because of the specific motoneuron adaptations needed to influence RFD. The experimental findings and computer simulations demonstrated that the neural adaptations required for increasing maximal force generation (peak motoneuron discharge rate at constant force) are different from those that influence contraction speed (recruitment speed and very early peak discharge rate during increasing force).

